# Isolation, sequence, infectivity and replication kinetics of SARS-CoV-2

**DOI:** 10.1101/2020.04.11.037382

**Authors:** Arinjay Banerjee, Jalees A. Nasir, Patrick Budylowski, Lily Yip, Patryk Aftanas, Natasha Christie, Ayoob Ghalami, Kaushal Baid, Amogelang R. Raphenya, Jeremy A. Hirota, Matthew S. Miller, Allison J. McGeer, Mario Ostrowski, Robert A. Kozak, Andrew G. McArthur, Karen Mossman, Samira Mubareka

## Abstract

SARS-CoV-2 emerged in December 2019 in Wuhan, China and has since infected over 1.5 million people, of which over 107,000 have died. As SARS-CoV-2 spreads across the planet, speculations remain about the range of human cells that can be infected by SARS-CoV-2. In this study, we report the isolation of SARS-CoV-2 from two cases of COVID-19 in Toronto, Canada. We determined the genomic sequences of the two isolates and identified single nucleotide changes in representative populations of our virus stocks. More importantly, we tested a wide range of human immune cells for productive infection with SARS-CoV-2. Here we confirm that human primary peripheral blood mononuclear cells (PBMCs) are not permissive for SARS-CoV-2. As SARS-CoV-2 continues to spread globally, it is essential to monitor single nucleotide polymorphisms in the virus and to continue to isolate circulating viruses to determine viral genotype and phenotype using *in vitro* and *in vivo* infection models.

---

A new coronavirus, severe acute respiratory syndrome coronavirus 2 (SARS-CoV-2) emerged in December 2019 in Wuhan, China (1). SARS-CoV-2 has since spread to 185 countries and infected over 1.7 million people, of which over 107,000 have died (2). The first case of COVID-19 was detected in Toronto, Canada on January 23, 2020 (3). Multiple cases have since been identified across Canada. As SARS-CoV-2 spreads globally, the virus is likely to adapt and evolve. It is critical to isolate SARS-CoV-2 viruses to characterize their ability to infect and replicate in multiple human cell types and to determine if the virus is evolving in its ability to infect human cells and cause severe disease. Isolating the virus also provides us with the opportunity to share the virus with other researchers to facilitate the development and testing of diagnostics, drugs and vaccines.

In this study, we describe how we isolated and determined the genomic sequence of SARS-CoV-2 from two cases of COVID-19 (SARS-CoV-2/SB2 and SARS-CoV-2/SB3-TYAGNC). In addition, we studied the replication kinetics of SARS-CoV-2/SB3-TYAGNC in human fibroblast, epithelial and immune cells. Importantly, we report that although a human lung cell line supported SARS-CoV-2 replication, the virus did not propagate in any of the tested immune cell lines or primary human immune cells. Interestingly, although we did not observe a productive infection in CD4^+^ primary T cells, we observed virus-like particles in these cells by electron microscopy. Our data shed light on a wider range of human cells that may or may not be permissive for SARS-CoV-2 replication and our studies strongly suggest that the human immune cells tested do not support a productive infection with SARS-CoV-2. Further studies are required to understand viral-host dynamics in immune cells.

## METHODS

### Cells

Vero E6 cells (African green monkey cells; ATCC, https://www.atcc.org) were maintained in Dulbecco’s modified Eagle’s media (DMEM) supplemented with 10% fetal bovine serum (FBS; Sigma-Aldrich, https://www.sigmaaldrich.com), 1x L-Glutamine and Penicillin/Streptomycin (Pen/Strep; Corning, https://ca.vwr.com). Calu-3 cells (human lung adenocarcinoma derived; ATCC, https://www.atcc.org) were cultured as previously mentioned (4). THF cells (human telomerase life-extended cells) were cultured as previously mentioned (5). THP-1 cells (monocytes; ATCC, https://www.atcc.org) were cultured in RPMI media (Gibco, https://www.thermofisher.com) supplemented with 10% FBS, 2mM L-glutamine, 1x Penicillin/streptomycin and 0.05 mM beta-mercaptoethanol. THP-1 cells were differentiated into macrophages and dendritic cells using 50 ng/ml GM-CSF (R&D Systems, https://www.rndsystems.com) + 50 ng/ml M-CSF (R&D Systems, https://www.rndsystems.com) and 50 ng/ml GM-CSF + 500 U/ml IL-4 (Biolegend, https://www.biolegend.com), respectively. PBMCs from healthy donors (n=2; OM8066 and OM8067) were purified into CD4^+^, CD8^+^, CD19^+^, monocytes and others (CD4^−^ CD8^−^ CD19^−^ cells) using CD4 negative selection kit (STEMCELL Technologies, https://www.stemcell.com), CD8 positive selection kit (STEMCELL Technologies, https://www.stemcell.com), PE positive selection kit (STEMCELL Technologies, https://www.stemcell.com) and Monocyte negative selection kit (STEMCELL Technologies, https://www.stemcell.com), respectively. For purity of cell types, see supplementary figure 1. CD4^+^, CD8^+^, CD19^+^ and CD4^−^ CD8^−^ CD19^−^ cells were resuspended in R-10 media (RPMI + 2 mM L-glutamine + 10% FBS + Penicillin/streptomycin) + 20 U/ml IL-2 (Biolegend, https://www.biolegend.com). Primary monocytes were resuspended in R-10 media. This work was approved by the Sunnybrook Research Institute Research Ethics Board (149-1994) and the Research Ethics Boards of St. Michael’s Hospital and the University of Toronto (REB 20-044; for PBMCs).

**Figure 1.**
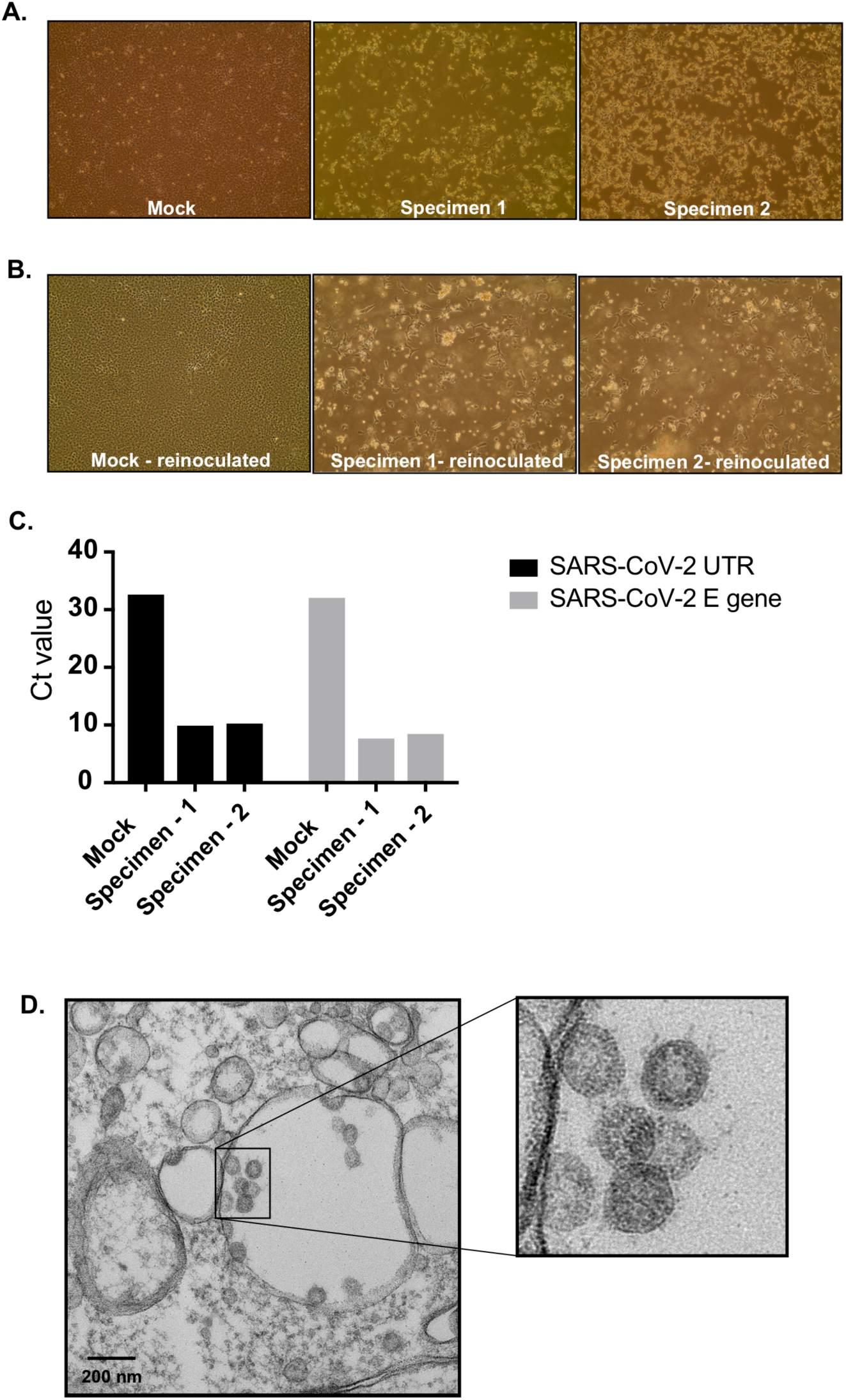
Isolating SARS-CoV-2 from COVID-19 patients. (A) Vero E6 cells were mock inoculated or inoculated with mid-turbinate clinical specimens from COVID-19 patients. Cells were incubated for 72 hours and observed for cytopathic effect (CPE) under a light microscope. (B) To determine if supernatant from Vero E6 cells that were mock inoculated or inoculated with clinical specimens contained replication competent virus, we re-inoculated fresh monolayer of Vero E6 cells and observed cells under a light microscope for CPE after 24 hours. (C) Quantitative real time PCR to detect SARS-CoV-2 5’-untranslated region (UTR) and envelope (E) gene in RNA extracted from supernatant that was collected from Vero E6 cells that were mock infected or infected with COVID-19 clinical specimens for 72 hours. (D) Electron micrograph of Vero E6 cells that were re-infected for 48 hours with supernatant that was collected from Vero E6 cells infected with clinical specimens. Insert – zoomed image of coronavirus-like particles.

### Virus isolation and quantification

Vero E6 cells were seeded at a concentration of 3×10^5^ cells/well in a 6-well plate. Next day, 200 μl of mid-turbinate swabs, collected from two COVID-19 patients was mixed with 200 μl of DMEM containing 16 μg/ml TPCK-treated trypsin and cells were inoculated. After 1 hr, the inoculum was replaced with DMEM containing 2% FBS and 6 μg/ml TPCK-treated trypsin. The cells were observed daily under a light microscope. Supernatant from the cells were used to determine virus titres (TCID_50_/ml) using Spearman and Karber’s method (6, 7) as outlined previously (8).

### Quantitative real time PCR

To detect SARS-CoV-2 in cell culture supernatant, 140 uL of supernatant was removed and detection of viral nucleic acids was performed by RT-PCR using an adaptation of Corman *et al.’s* protocol (9). Briefly, viral RNA was extracted from infected cells using QIAamp viral RNA kit (Qiagen, https://www.qiagen.com) according to manufacturer’s instructions. The RT-PCR reactions were carried out using Luna Universal qPCR Master Mix (New England Biolabs, https://www.neb.ca) according to manufacturer’s instructions. Two separate gene targets were used for detection, the 5’ untranslated region (UTR) and the envelope (E) gene. Primers and probes used were: 5’-UTR For: GTTGCAGCCGATCATCAGC; 5’-UTR Rev: GACAAGGCTCTCCATCTTACC; 5’-UTR probe: FAM-CGGTCACACCCGGACGAAACCTAG-BHQ-1; E-gene For: ACAGGTACGTTAATAGTTAATAGCGT; E-gene Rev: ATATTGCAGCAGTACGCACACA; E-gene probe: CAL Fluor Orange 560-ACACTAGCCATCCTTACTGCGCTTCG-BHQ-1. The cycling conditions were: 1 cycle of denaturation at 60°C for 10 minutes then 95°C for 2 minutes followed by 44 amplification cycles of 95°C for 10s and 60°C for 15s. Analysis was performed using the Rotor-Gene Q software (Qiagen, https://www.qiagen.com) to determine cycle thresholds (Ct).

### Electron Microscopy

Samples were fixed in 10% neutral buffered formalin (Sigma-Aldrich, https://www.sigmaaldrich.com/) for 1 hr. Pellets were washed with 0.1 M phosphate buffer (pH 7.0) and post-fixed with 1% osmium tetroxide in 0.1 M phosphate buffer (pH 7.0) for 1 hr. Pellets were washed with distilled water and en-bloc stained with 2% uranyl acetate in distilled water for 2 hrs. Pellets were washed with distilled water and dehydrated in an ethanol series. Pellets were infiltrated with Embed 812/Araldite resin and cured at 65 degrees for 48 hrs. Resin blocks were trimmed, polished and 90 nm thin sections were ultramicrotomed (Leica Reichert Ultracut E, https://www.leica-microsystems.com) and mounted on TEM grids. Thin sections were stained with 5% uranyl acetate and 5% lead citrate. Sections were imaged using Transmission Electron Microscopy (ThermoFisher Scientific Talos L120C, https://www.thermofisher.com) using a LaB6 filament at 120kV. 10 fields per cell type were scanned. Each field was imaged with different magnifications at 2600x, 8500x, 17500x and 36000x.

### Flowcytometry

To prepare cells for flowcytometry, 100μL (400,000 cells) of primary CD4^+^, CD8^+^, CD19^+^ and monocytes were washed with 1 ml of phosphate buffered saline (PBS) and spun at 500 g for 5 minutes. The cells were resuspended in 100 μL of Live/Dead Violet (ThermoFisher Scientific, https://www.thermofisher.com) as per manufacturer’s recommendation and diluted 1:1000 in PBS. Cells were incubated at 4°C for 30 minutes. Next, cells were washed with 1 ml of FACS buffer (in-house reagent) and spun at 500 g for 5 minutes. Cells were then stained with 100 μL of their respective stains (*α*CD4-FITC, *α*CD8-FITC, *α*CD19-FITC, *α*CD14-APC; Biolegend, https://www.biolegend.com) at a concentration of 1μg/mL for 30 min at 4°C. After staining, the cells were washed with 1mL of FACS Buffer and spun at 500 g for 5 minutes. Extra aliquots of cells were left unstained, which were also spun at 500g for 5 minutes. The pellets were resuspended in 100 μL of 1% paraformaldehyde (PFA; ThermoFisher Scientific, https://www.thermofisher.com) and analyzed. Samples were run on the BD LSR Fortessa X20 (BD, https://www.bdbiosciences.com). Cells were gated on Live/Dead negative to exclude debris and dead cells and were then gated on their respective cell surface markers to assess purity.

### Sequencing

RNA was extracted from the supernatant of Vero E6 cells after 1 passage using the QIAamp Viral RNA Mini kit (Qiagen, https://www.qiagen.com/us/) without addition of carrier RNA. dsDNA for sequencing library preparation was synthesized using the Liverpool SARS-CoV-2 amplification protocol (10). Two 100 µM primer pools were prepared by combining primer pairs in an alternating fashion to prevent amplification of overlapping regions in a single reaction. In a PCR tube, 1µL Random Primer Mix (ProtoScript II First Strand cDNA Synthesis Kit, New England Biolabs, https://www.neb.ca) was added to 7 µL extracted RNA and denatured on SimpliAmp thermal cycler (ThermoFisher Scientific, https://www.thermofisher.com) at 65°C for 5 min and then incubated on ice. 10 µL 2X ProtoScript II Reaction Mix and 2 µL 10X ProtoScript II Enzyme Mix were then added to the denatured sample and cDNA synthesis performed using the following conditions: 25°C for 5 min, 48°C for 15 min and 80°C for 5 min. After cDNA synthesis, in a new PCR tube 2.5 µL cDNA was combined with 12.5 µL Q5 High-Fidelity 2X Master Mix (NEB, Ipswich, USA), 8.8 µL nuclease free water (ThermoFisher Scientific, https://www.thermofisher.com), and 1.125 µL of 100 µM primer pool #1 or #2. PCR cycling was then performed as follows: 98°C for 30 sec followed by 40 cycles of 98°C for 15 sec and 65°C for 5 min.

All PCR reactions were purified using RNAClean XP (Beckman Coulter, https://www.beckmancoulter.com) at 1.8x bead to amplicon ratio and eluted in 30 µL. 2 µL of amplified material was quantified using a Qubit 1X dsDNA (ThermoFisher Scientific, https://www.thermofisher.com) following the manufacturer’s instructions. Illumina sequencing libraries were prepared using Nextera DNA Flex Library Prep Kit and Nextera DNA CD Indexes (Illumina, https://www.illumina.com) according to manufacturer’s instructions. Paired-end 150 bp sequencing was performed for each library on a MiniSeq with the 300-cycle mid-output reagent kit (Illumina, https://www.illumina.com), multiplexed with targeted sampling of ∼40,000 clusters per library. Sequencing reads from pool #1 and pool #2 were combined (as R1 and R2), amplification primer sequences removed using cutadapt (version 1.18) (11), and Illumina adapter sequences were removed and low quality sequences trimmed or removed using Trimmomatic (version 0.36) (12). Final sequence quality and confirmation of adapter/primer trimming were confirmed by FASTQC (version 0.11.5) (13). SARS-CoV-2 genome sequences were assembled using UniCycler (version 0.4.8; default settings, except for --mode conservative) (14) and assembly statistics generated by QUAST (version 5.0.2) (15). Sequencing depth and completeness of coverage of the assembled genomes was additionally assessed by Bowtie2 (version 2.3.4.1) (16) alignment of the sequencing reads against the assembled contigs and statistics generated by ngsCAT (version 0.1) (17). Sequence variation in the assembled genomes was assessed by BLASTN against SARS-CoV-2 genome sequences available in GenBank as well as BreSeq (version 0.35.0) (18) analysis relative to GenBank entry MN908947.3 (first genome sequence reported from the original Wuhan outbreak, China).

## RESULTS

For virus isolation, we inoculated Vero E6 cells with the samples collected from mid-turbinate swabs and monitored for cytopathic effects (CPE) daily. Seventy-two hours post infection (hpi), cells inoculated with both samples displayed extensive CPE, relative to mock-inoculated cells (Figure 1, panel A). We collected 200 uL of cell culture supernatant and re-infected a fresh layer of Vero E6 cells. Twenty-four hours post infection, both wells containing cells that were re-inoculated displayed extensive CPE (Figure 1, panel B). We extracted viral RNA from the supernatant and confirmed the presence of SARS-CoV-2 using a diagnostic real-time PCR assay (qPCR; Figure 1, panel C). We also confirmed the presence of coronavirus-like particles in infected Vero E6 cells by electron microscopy (Figure 1, panel D).

Next, we performed genome sequencing of both isolates, generating genome sequences with 7500-8000 fold coverage and ∼94% completeness, with only ∼260 bp and ∼200 bp at the 5’ and 3’ termini undetermined (Table 1). Both shared synonymous and non-synonymous substitutions with those independently observed in direct sequencing of clinical isolates (Table 1; Mubareka & McArthur, unpublished). SARS-CoV-2/SB2 additionally contained a non-synonymous substitution at position 2832 (K856R in ORF1ab polyprotein) and three regions with mutations or a deletion supported by a minority of sequencing reads, while SARS-CoV-2/SB3-TYAGNC only had an additional synonymous substitution in ORF1ab polyprotein (Y925Y) plus a minority of sequencing reads supporting another synonymous substitution in the ORF3a protein (I7I). As such, SARS-CoV-2/SB3-TYAGNC was used for subsequent studies as best representative of a clinical viral isolate. Raw sequencing reads for each isolate are available in NCBI BioProject PRJNA624792. Only sequencing reads that aligned by Bowtie2 to the MN908947.3 SARS-CoV-2 genome were included in the deposited sequence files.

**Table 1.**
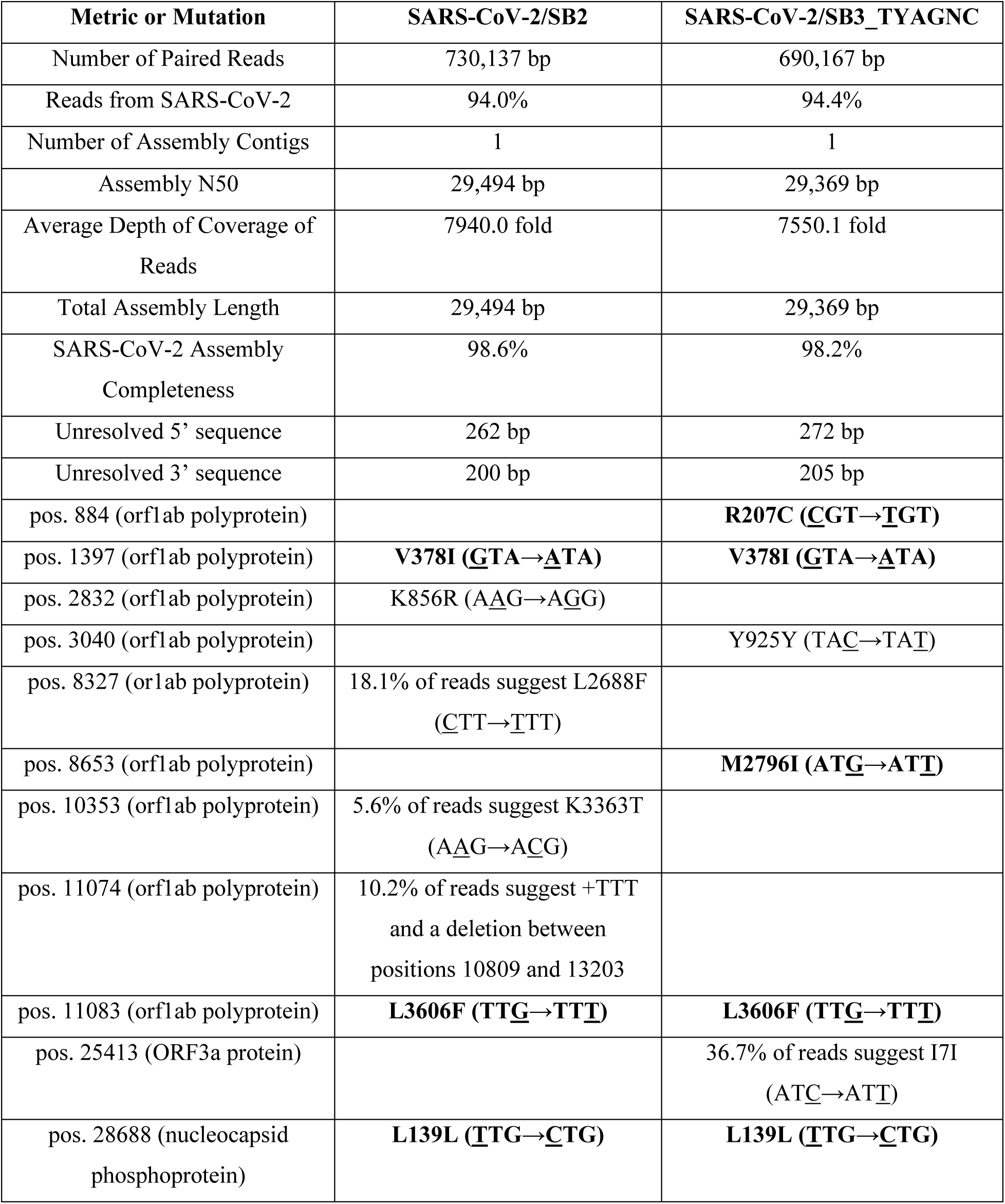
Sequencing read and genome assembly statistics. Predicted mutations are relative to the MN908947.3 SARS-CoV-2 genome (29,903 bp). Mutations within codons are underlined. All mutations were predicted by 100% of sequencing reads mapping to that position, unless otherwise noted. None of the mutations with support from less than 100% of sequencing reads appeared in the final assembled genome consensus sequences. Substitutions in bold have been observed in direct sequencing of patient isolates (Mubareka & McArthur, unpublished).

To determine the replication kinetics of SARS-CoV-2 in human structural and immune cells, we infected Calu-3 (lung adenocarcinoma epithelial), THF (telomerase life extended human fibroblasts), Vero E6 (African green monkey kidney epithelial), THP-1 (monocytes and differentiated macrophages and dendritic cells) and primary peripheral blood mononuclear cells (PBMCs) from healthy human donors (CD4^+^, CD8^+^, CD19^+^, monocytes and other PBMCs; Supplementary Figure S1) with a multiplicity of infection (MOI) of 0.01. We monitored virus replication over a period of 72 hours in the cell lines (Figure 2). We also determined virus replication in PBMCs from healthy donors over a period of 48 hours (Figure 2). SARS-CoV-2 propagated to high titres in Vero E6 and Calu-3 cells (Figure 2). SARS-CoV-2 did not replicate efficiently in THF cells (Figure 2). Interestingly, human immune cell lines and primary PBMCs from healthy donors did not support SARS-CoV-2 replication (Figure 2)

**Figure 2.**
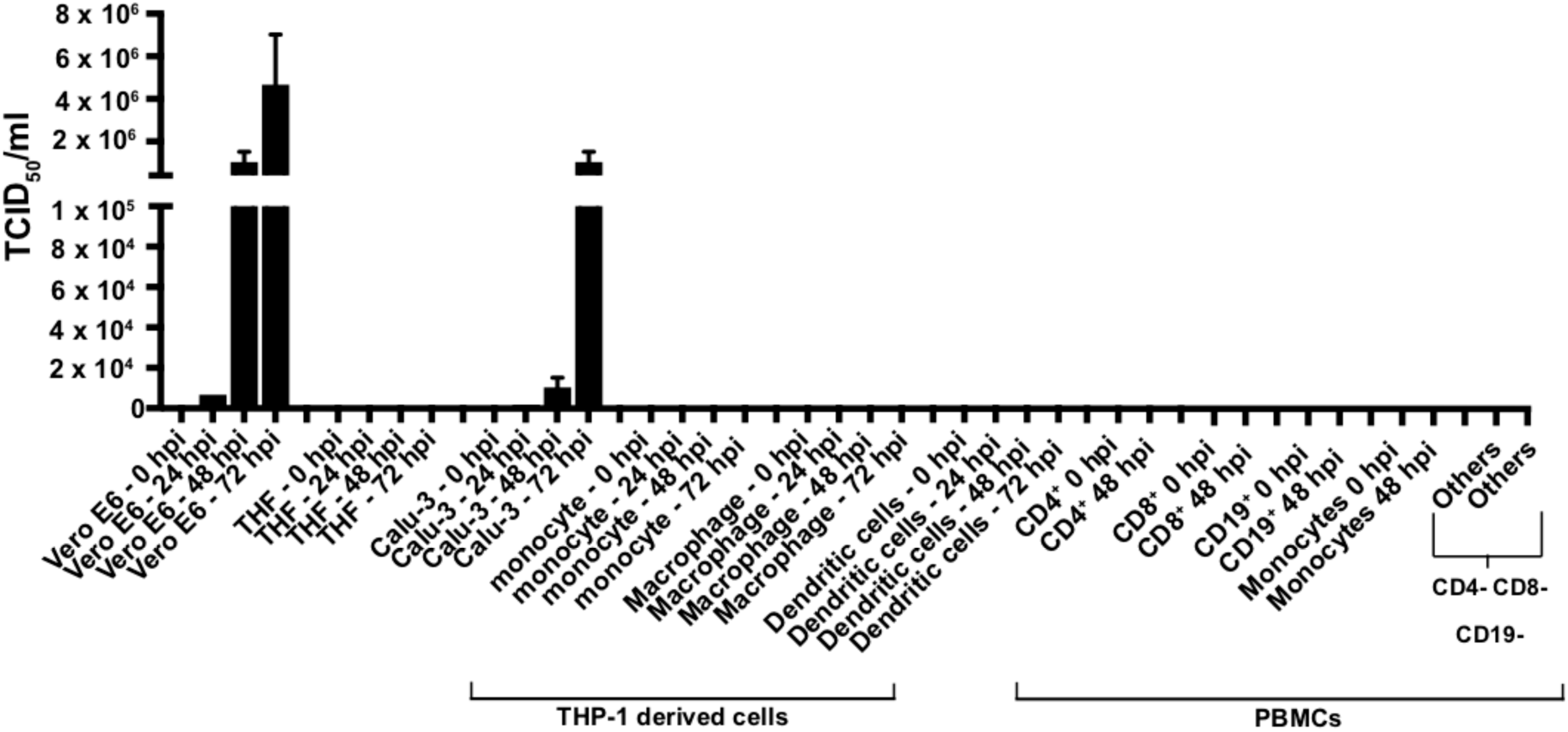
Replication of SARS-CoV-2 in human structural and immune cells. To determine human cells that support SARS-CoV-2 replication, we infected human cell lines and primary cells at an MOI of 0.01 (n = 2 independent experiments; supernatant from each experiment was titrated in triplicates). We infected Vero E6 cells as a control. THF and Calu-3 cells represent human structural cells. THP-1 is a monocyte cell line that was used to derive macrophages and dendritic cells. PBMCs from healthy human donors (n = 2 independent donors) were used to generate CD4^+^, CD8^+^, CD19^+^, monocytes and other (CD4^−^ CD8^−^ CD19^−^) cell populations. Supernatant from infected cells were collected at various times and titrated on Vero E6 cells to determine virus titres (TCID_50_/ml). hpi, hours post infection; CD, cluster of differentiation; PBMC, peripheral blood mononuclear cells.

To further support virus replication data, we imaged infected human epithelial, fibroblast and immune cells using electron microscopy after 48 hrs of infection with SARS-CoV-2 at a MOI of 0.01 (Figure 3). We scanned 10 different fields per cell type, each using four different magnifications of 2600x, 8500x, 17500x and 36000x to determine if the cell populations contained virus-like particles. Virus-like particles were detected in 7/10 fields in Vero E6 cells and 8/10 fields in Calu-3 cells (Figure 3, panels A and B). Interestingly, we also detected virus-like particles in 2/10 fields in primary CD4^+^ T cells (Figure 3, panel C). We did not observe any virus-like particles in other human immune cells that were experimentally infected with SARS-CoV-2 (Figure 3, panels D-J).

**Figure 3.**
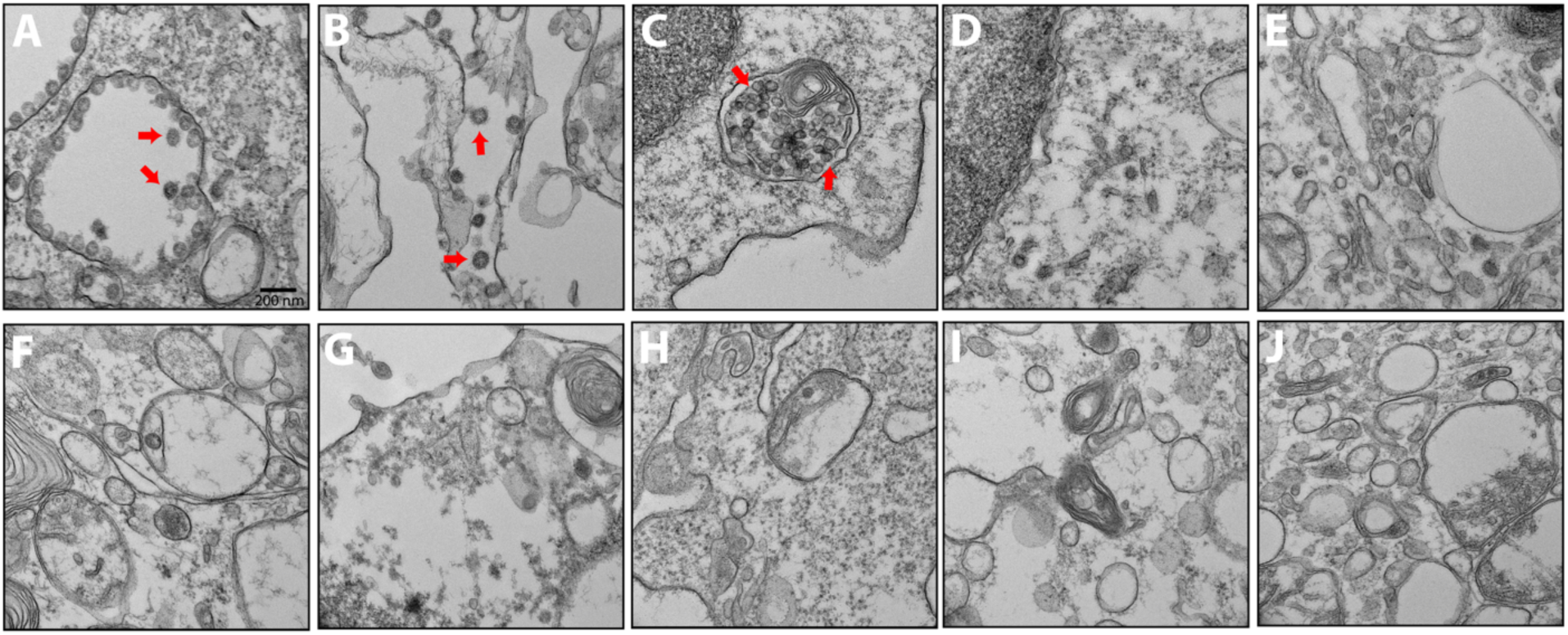
Electron micrographs of SARS-CoV-2 infected cells. To detect coronavirus-like particles in experimentally infected human structural and immune cells, we infected a range of cells with SARS-CoV-2 at a MOI of 0.01 for 48 hours. The cells were fixed, processed and imaged using a transmission electron microscope (n = 10 fields / cell type). Representative image of each cell line is shown above. Virus-like particles are indicated by red arrows. (A) Vero cells. (B) Calu-3 cells. (C) CD4^+^ PBMC. (D) CD8^+^ PBMC. (E) CD19^+^ PBMC. (F) Monocytes from PBMC. (G) Other cells from PBMC (CD4^−^ CD8^−^ CD19^−^ cell populations). (H) THP-1 monocyte. (I) THP-1-derived macrophage. (J) THP-1-derived dendritic cell.

## DISCUSSION

In this study, we report the isolation of two replication competent SARS-CoV-2 isolates from COVID-19 patients in Canada. We used TPCK-treated trypsin to enhance infection using clinical specimens (Figure 1, panel A). Exogenous trypsin activates SARS-CoV spike proteins more efficiently and facilitates cellular entry (19). Exogenous trypsin treatment has also been shown to enhance infectivity of other zoonotic bat-borne coronaviruses (20). Furthermore, TPCK-treated trypsin was used to successfully isolate SARS-CoV-2 in China (1). In our studies, subsequent infection and virus replication did not require any additional TPCK-treated trypsin (Figure 1, panel B). The presence of CPE alone does not indicate the successful isolation of a coronavirus. Mid-turbinate samples from adults with acute respiratory distress may often contain other microbes, including viruses (21). Thus, to identify our isolates, we sequenced them to confirm that they were reflective of SARS-CoV-2 isolates infecting patients worldwide, selecting SARS-CoV-2/SB3-TYAGNC for experimental investigation since this isolate produced less minority sequencing reads (Table 1).

SARS-CoV caused the 2003-04 outbreak of SARS. SARS-CoV can infect structural (22) and immune cell lines (23) from humans *in vitro*. We infected a range of human cell populations with SARS-CoV-2/SB3-TYAGNC to identify cell types that can support productive infection of SARS-CoV-2. Both Vero E6 and Calu-3 cells supported SARS-CoV-2 replication to high titres (Figure 2), as reported in other recent studies (24, 25). Previously, SARS-CoV was also shown to replicate efficiently in Vero E6 cells (22). Vero E6 cells are immunodeficient, with deficiencies in innate antiviral interferon signaling, which makes them ideal candidates for virus isolation (26). However, to enable studies on SARS-CoV-2-host interactions, it is important to identify human lung epithelial cells with intact immune responses that can support SARS-CoV-2 replication. We and others have previously shown that SARS-CoV and Middle East respiratory syndrome coronavirus (MERS-CoV) replicate efficiently in Calu-3 cells (8, 27, 28). In addition, SARS-CoV- and MERS-CoV-induced immune responses have been studied in Calu-3 cells (28, 29). The ability to infect Calu-3 cells with SARS-CoV-2 (Figure 2) will facilitate *in vitro* studies into virus-host interactions using SARS-CoV-2. Other commonly used human lung cells, such as A549 do not support efficient replication of SARS-CoV-2 (24). Furthermore, hTERT life-extended human fibroblast cells (THF) also did not support virus replication (Figure 2).

Previous studies have shown that human immune cells, such as THP-1 cell lines are susceptible to SARS-CoV infection (23). In our study, human immune cell populations, including THP-1-derived cell lines and primary cells (PBMCs) did not support productive SARS-CoV-2 replication (Figure 2). Interestingly, although primary CD4^+^ T cells did not support productive virus replication, we observed virus-like particles in these cells by electron microscopy (Figure 3, panel C). This is consistent with a recent report by Wang and colleagues where they demonstrated that human T cell lines are susceptible to SARS-CoV-2 and pesudotyped viruses (30). However, the study by Wang *et al.* did not quantify virus titres in the supernatant from infected cells. In our study, we could not detect any replication competent virus in the supernatant that was collected from SARS-CoV-2 infected CD4^+^ T cells (Figure 3). Human immune cells lack expression of angiotensin converting enzyme 2 (ACE2) (31) (https://www.proteinatlas.org), the functional receptor of SARS-CoV-2 (1, 32). Thus, although it is intriguing that CD4^+^ T cells may be susceptible to SARS-CoV-2, our data show that these cells are not permissive to SARS-CoV-2 replication *in vitro*.

In this study, we show that primary human T cells (CD4^+^ and CD8^+^) do not support productive virus replication. However, our electron micrographs demonstrate that SARS-CoV-2 likely replicates in CD4^+^ T cells, but the replication cycle is likely terminated prior to virus maturation and egress. Thus, more work is needed to fully identify the susceptibility and permissivity of CD4^+^ T cells to SARS-CoV-2.

## ACKNOWLEDGEMENTS

We would like to acknowledge Lindsey Fiddes’ help with electron microscopy. SARS-CoV-2 Liverpool protocol genome amplification primer sequences were generously shared by Public Health England.

This study was supported by a Canadian Institutes of Health Research (CIHR) COVID-19 rapid response grant to principal applicant K.M. and Co-Applicants A.B., A.G.M., M.S.M. and S.M. A.B. is funded by the Natural Sciences and Engineering Research Council of Canada (NSERC). J.A.N. was supported by funds from the Comprehensive Antibiotic Resistance Database. B.P.A. and A.R.R. were supported by Canadian Institutes of Health Research (CIHR) funding (PJT-156214 to A.G.M.). Computer resources were supplied by the McMaster Service Lab and Repository computing cluster, funded in part by grants to A.G.M. from the Canadian Foundation for Innovation. Additional cloud computing needs were funded by the Comprehensive Antibiotic Resistance Database. J.A.H. is supported by the Canada Research Chairs Program and an Ontario Early Career Researcher Award. M.S.M. is supported by a CIHR COVID-19 rapid response grant, a CIHR New Investigator Award and an Ontario Early Researcher Award.

