## supplementary figure 1 for "Isolation, sequence, infectivity and replication kinetics of SARS-CoV-2"

**Title:** Sequence, infectivity and replication kinetics of SARS-CoV-2 isolated from COVID-19 patients in Canada

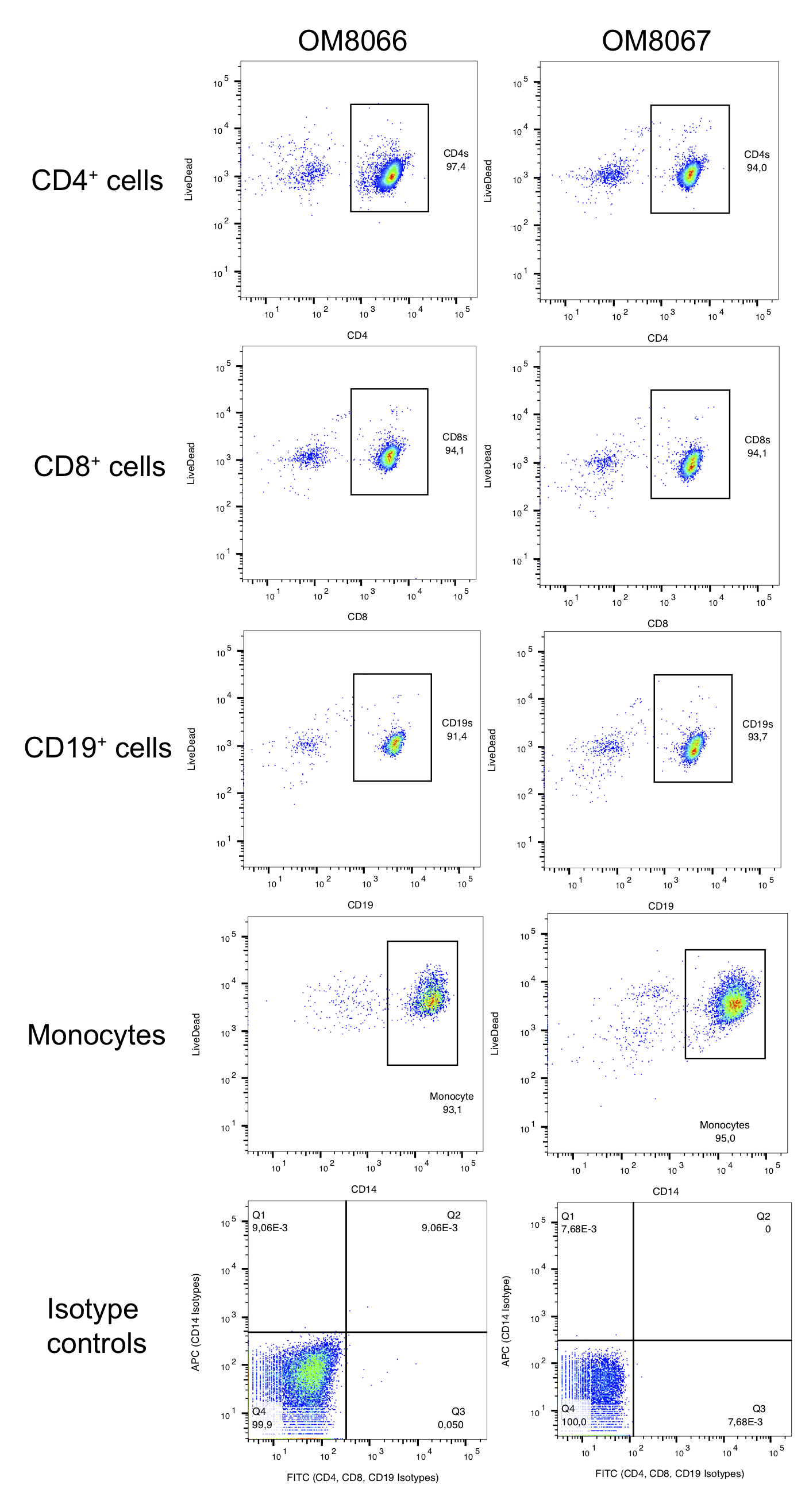

**Figure S1. Percentage purities of cell populations that were purified from human PBMCs.** Human PBMCs were collected (n = 2 independent healthy donors) and purified using cell-type specific purification kits (see Methods). Cells were stained for their respective markers and analyzed using flowcytometry. Percentage purity of each cell population are mentioned in the respective panels.
